# Structure, culture, and predicted function of the gut microbiome of the Mormon cricket *Anabrus simplex* (Orthoptera: Tettigoniidae)

**DOI:** 10.1101/080846

**Authors:** Chad C. Smith, Robert B. Srygley, Frank Healy, Karthikeyan Swaminath, Ulrich G. Mueller

## Abstract

1. The gut microbiome of insects plays an important role in their ecology and evolution, participating in nutrient acquisition, immunity, and behavior. Microbial community structure within the gut is heavily influenced by differences among gut regions in morphology and physiology, which determine the niches available for microbes to colonize.
2. We present a high-resolution analysis of the structure of the gut microbiome in the Mormon cricket *Anabrus simplex,* an insect known for its periodic outbreaks in the western United States and nutrition-dependent mating system. The Mormon cricket microbiome was dominated by eleven taxa from the Lactobacillaceae, Enterobacteriaceae, and Streptococcaeae. While most of these were represented in all gut regions, there were marked differences in their relative abundance, with lactic-acid bacteria (Lactobacillaceae) more common in the foregut and midgut and enteric (Enterobacteriaceae) bacteria more common in the hindgut.
3. Differences in community structure were driven by variation in the relative prevalence of three groups: a *Lactobacillus* in the foregut, *Pediococcus* lactic-acid bacteria in the midgut, and *Pantoea agglomerans*, an enteric bacterium, in the hindgut. These taxa have been shown to have beneficial effects on their hosts in insects and other animals by improving nutrition, increasing resistance to pathogens, and modulating social behavior.
4. Using PICRUSt to predict gene content from our 16S rRNA sequences, we found enzymes that participate in carbohydrate metabolism and pathogen defense in other orthopterans. These were predominately represented in the hindgut and midgut, the most important sites for nutrition and pathogen defense.
5. Phylogenetic analysis of 16S rRNA sequences from cultured isolates indicated low levels of divergence from sequences derived from plants and other insects, suggesting that these bacteria are likely to be exchanged between Mormon crickets and the environment.
6. Our study shows strong spatial variation in microbiome community structure, which influences predicted gene content and thus the potential of the microbiome to influence host function.

## INTRODUCTION

Insects are the most speciose and abundant taxa in the animal kingdom, playing a key ecological role in many of the world’s ecosystems. Symbioses between insects and their microbial associates has undoubtedly contributed to their success, providing the capability to degrade recalcitrant food, to supplement nutrient-deficient diets, to protect them from their natural enemies, and to modulate the expression of social behavior (Engel & Moran 2013; Douglas 2015). Among the niches available to occupy the host, the gut houses the largest and most diverse microbiome in insects (Engel & Moran 2013; Douglas 2015) and other animals (Ley et al. 2008; Cho & Blaser 2012). Gut morphology and physiology vary markedly along the alimentary tract, resulting in an environmental gradient that influences, and is influenced by, the microbial communities that populate it (Dillon & Dillon 2004; Engel & Moran 2013).

The insect gut consists of three regions that are analogous to that in mammals, the foregut, the midgut and the hindgut, each of which contributes to a different aspect of gut function (Douglas 2013). The foregut serves as the entry point for food, where it is stored in the crop before passing through the proventriculus, a valve that can also be modified to mechanically filter food (Woodring & Lorenz 2007; Douglas 2013) and even microbes (Lanan et al. 2016). Digestion and absorption of nutrients begins at the midgut, which, in some species, contains specialized crypts that house microbes that aid in insect nutrition (Kikuchi, Meng & Fukatsu 2005; Bistolas et al. 2014). Host immune factors have been shown to play an important role in regulation of commensal microbes in the midgut (Ryu, Ha & Lee 2010; Buchon, Broderick & Lemaitre 2013), some of which protect the host from pathogens (Forsgren et al. 2010). Following the midgut is the hindgut, which is comprised of the ileum, colon, and rectum. Malphigian tubules permeate the anterior hindgut, excreting nitrogenous waste and other solutes from the hemocoel that can provide nutrients for dense populations of microbes (Bignell 1984). In some species, dense bristle-like structures in the ileum (Woodring & Lorenz 2007) and rectal papillae (Hunt & Charnley 1981) provide attachment sites for bacteria, some of which fix nitrogen (Tai et al. 2016), degrade recalcitrant plant polymers (Kaufman & Klug 1991; Engel & Moran 2013), and prevent infection (Dillon & Charnley 2002).

The Mormon crickets *Anabrus simplex* (Orthoptera: Tettigoniidae) is an economically important shield-backed katydid distributed throughout the Western United States. Mormon crickets can form dense aggregations of millions of individuals spread over 10 kilometers long and several kilometers wide, feeding on forbes, grasses, and agricultural crops as they march in migratory bands across the landscape (MacVean 1987; Simpson et al. 2006). Mormon crickets are also emerging as a model for the study of how social interactions and diet influence immunity (Srygley et al. 2009; Srygley & Lorch 2011) and the microbiome (Smith et al. 2016). Differences in population density are linked to reproductive behavior, as in high density populations, protein-limited females compete for access to males to gain access to a proteinaceous “nuptial gift” males produce for females during copulation (Gwynne 1984). While consumption of male nuptial gifts by females does not influence the composition of the microbiome, sexually inactive females experience a dramatic decline in *Pediococcus* lactic-acid gut bacteria compared to sexually active females (Smith et al. 2016). Lactic-acid bacteria are common associates of the alimentary tract and regarded for their beneficial effects on immune function and nutrition in animals, including insects (Forsgren et al. 2010; Storelli et al. 2011; Erkosar et al. 2015).

We characterize the structure of the gut microbiome of Mormon crickets and infer their evolutionary relationships using a combination of culture-dependent and culture-independent approaches. Our aims are to determine whether gut microbial communities vary along the alimentary tract in the Mormon cricket and to infer their potential to influence host function based on their known taxonomic associations with other insects and by employing bioinformatic tools that predict metabolic capabilities from 16S rRNA sequences. We also establish methods for isolating Mormon cricket gut microbiota in culture to permit future experimental manipulations of the gut microbiome and build genomic resources to infer their evolution and function.

## MATERIALS AND METHODS

### Animal collection and tissue processing

Mormon crickets were obtained from field (n=5) and laboratory-raised (n=8) collections. Wild females were caught in EK Mountain (43°47’58”N, 106°50’31”W, 1752 m) near Kaycee, Wyoming in the summer of 2014, immediately preserved in 100% ethanol, and stored at −80°C until dissection. Laboratory-raised Mormon crickets were derived from eggs collected from individuals caught on Paint Rock Road (44°27'52"N, 107°27'37"W, 2654 m) in the Bighorn Mountains and fed a mixture of wheat bran, wheat germ, sunflower, mixed bird seeds, tropical fish flakes, fresh Romaine lettuce (added daily), and water *ad libitum*.

Mormon crickets were dissected using flame-sterilized tools after rinsing in 1% bleach for 3 min followed by two rinses in autoclaved distilled water to remove bacteria on the exoskeleton. DNA from the foregut (crop and proventriculus), midgut (ventriculus), ileum, and rectum (Fig. 1) of laboratory-raised crickets was extracted with MoBio Powersoil^©^ as in Smith et al. (2016). Foregut (crop and proventriculus), midgut (ventriculus), and hindgut tissue (ileum and rectum combined) was extracted from field-collected animals using a bead-beating/phenol-chloroform extraction protocol (see supplementary material). DNA extraction methods can influence the representation of taxa in 16S rRNA metagenomic studies (Yuan et al. 2012), however our aim here is not to make inferences about differences between field and laboratory-raised animals but differences among tissue types. We include the source of the animal (field or laboratory) and as a covariate in our statistical analyses to account for variation due to source/DNA extraction method (see Statistics).

**Figure 1.**
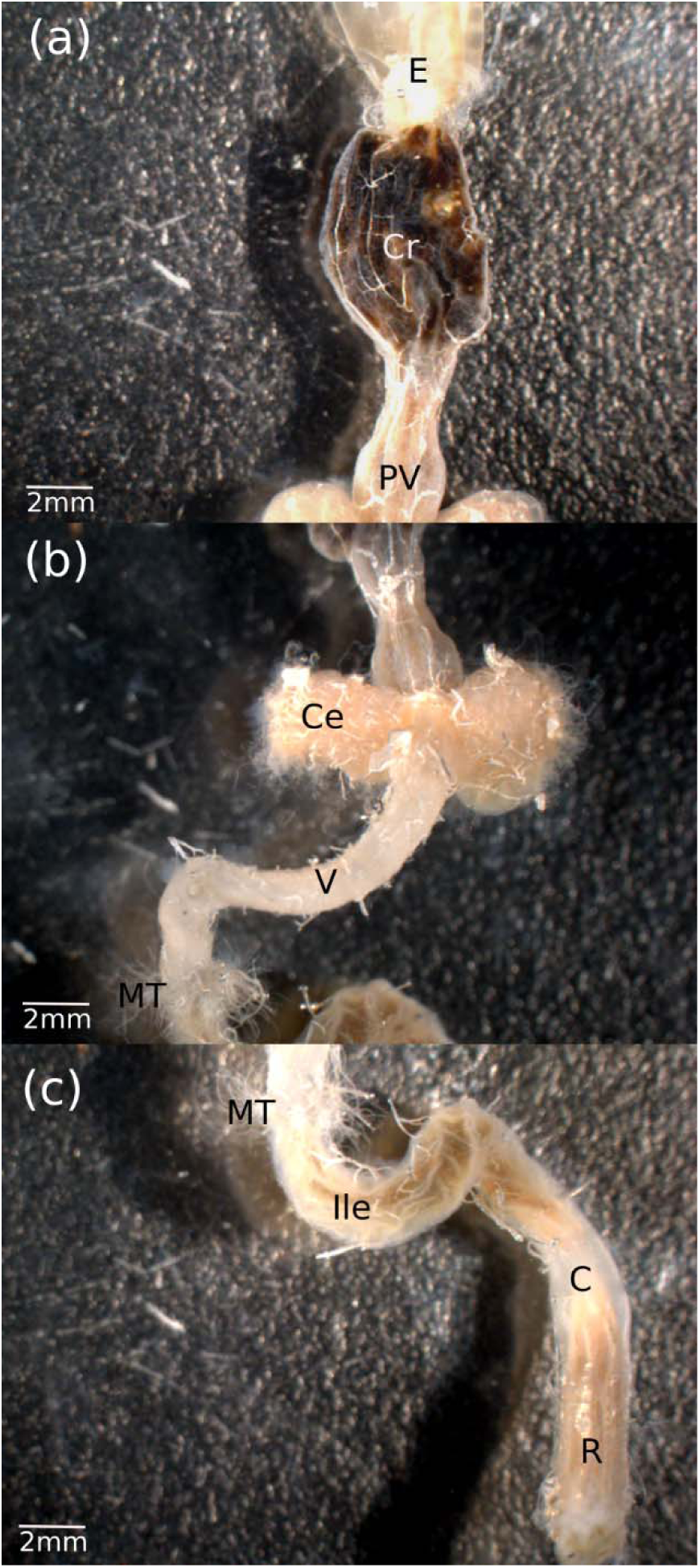
External morphology of the (a) foregut, (b) midgut, and (c) hindgut in the Mormon cricket. E=esophagus, Cr=crop, PV=proventriculus, Ce=cecum, V=ventriculus, MT=Malphigian tubules, Ile=ileum, C=colon, R=rectum. Malphigian tubules have been trimmed to illustrate their entry point into the hindgut.

### Sequencing and Bioinformatics

The variable V4 region of 16S rRNA gene was amplified with universal primers (Hyb515F: 5'-GTGYCAGCMGCCGCGGTA −3', Hyb806R: 5'-GGACTACHVGGGTWTCTAAT-3') and sequenced on the Illumina Miseq V3 platform. DADA2 1.1.5 (Callahan et al. 2016) was used to process the raw sequencing data and taxonomy was assigned with the Greengenes 13.8 database at 97% identity (see supplementary material). Sequence variants that comprised an average of less than 1% of the reads recovered within a given Mormon cricket were removed prior to analysis using phyloseq 1.16.2 (McMurdie & Holmes 2013).

### Metagenomic predictions

We used PICRUSt (Phylogenetic Investigation of Communities by Reconstruction of Unobserved States) v1.1.0 (Langille et al. 2013) to estimate the functional gene content of our samples. PICRUSt generates metagenomic predictions from 16S rRNA data using annotations of sequenced genomes in the IMG database. Nearest sequenced taxon index (NSTI) values were small (mean ± sd: 0.03 ± 0.01, range: 0.006-0.040), indicating the taxa in our samples were closely related to the genomes in the IMG database (see supplementary material). Greengenes IDs used by PICRUSt to construct the phylogenetic tree were assigned to sequence variants using Qiime 1.9 (Caporaso et al. 2010), and the Kyoto Encyclopedia of Genes and Genomes (KEGG) database was used for functional classification.

### Bacterial Abundance

The abundance of bacteria was estimated using qPCR following Powell et. al (2014) from laboratory-raised (n=8) and field-caught (n=8) Mormon crickets (see supplementary information). Universal 16S rRNA gene primers 27F (5’-AGAGTTTGATCCTGGCTCAG-3’) and 355R (5’-CTGCTGCCTCCCGTAGGAGT-3’) were used to amplify all copies of the 16S rRNA gene in tissue samples from laboratory (n=8) and field caught individuals (n=8) and copy number quantified with standard curves from the cloned target sequence (Powell et al. 2014)

### Culturing and phylogenetic analysis

Five lab-reared female Mormon crickets were surface sterilized in 1% bleach for three min, rinsed twice in sterile water and dissected using flame-sterilized tools. Gut tissue was homogenized for 10 s with a bead beater using autoclaved 3.2mm stainless steel beads in sterile PBS. Homogenates were plated onto trypsin soy agar, brain heart infusion agar, nutrient agar, or Man–Rogosa–Sharpe agar (BD), cultured in anaerobic or Campy (low O_2_) Gaspak pouches (Becton, Dickinson and Company, Franklin Lakes, NJ) at 37°C for 24-48 hours, and individual colonies passaged three times to obtain pure isolates. DNA was then extracted with chelex and the 16S gene amplified for Sanger sequencing using 27F (5’-AGAGTTTGATCCTGGCTCAG-3’) and 1492R (5’-GGTTACCTTGTTACGACTT-3’) primers (see supplementary material).

### Phenotypic assays

Fresh overnight cultures of all isolates were used for microscopic analysis. Lactobacillaceae isolates were cultured in Man–Rogosa–Sharpe medium and Enterobacteriaceae were cultured in nutrient broth or LB medium. Biochemical tests were done following Bridson (1998). Motility was determined using SIM medium and microscopic examination of culture wet mounts. Man–Rogosa–Sharpe or nutrient broth containing 1 g/L potassium nitrate was used for nitrate reduction tests. Fermentation tests were done anaerobically in Man–Rogosa–Sharpe and nutrient broth media with the addition of indicated sugars to 1% w/v final concentration.

### Statistics

Analyses were performed in R 3.3.1 (R Core Development Team 2013). Sequence tables were rarified at 1300 reads using phyloseq (McMurdie & Holmes 2013), resulting in the exclusion of hindgut samples from two field-caught females that had a low number of reads. Alpha diversity was compared among tissue types and between origin of subject (field vs. lab) with a linear mixed model (Bates et al. 2013), entering the individual ID as a random effect to account for within-subject correlations in diversity. Species richness, Chao1, and the Shannon-Weiner diversity index were calculated. Post-hoc comparisons among gut regions were performed using a Tukey test (Hothorn, Bretz & Westfall 2008).

Beta diversity among gut tissue types and between animal source (field vs. lab) was assessed with a distance-based redundancy analysis (db-RDA) in vegan 2.3 (Oksanen et al. 2015), specifying a principal components ordination of Bray-Curtis distances. Statistical significance of the terms in the db-RDA model was determined by 999 permutations of the distance matrix in vegan, restricting the permutations to within each individual to retain the nested structure of the data. The same procedure was also used to examine variation among tissue types in the abundance of KEGG pathways, except nonmetric multidimensional scaling was used for ordination.

We assessed the difference in taxon abundance among tissue types in univariate analyses by fitting the data to a negative binomial generalized linear mixed model (Bates et al. 2013), specifying the individual ID as the random effect and the tissue type and animal source (field vs. lab) as fixed effects. A similar procedure was used to assess differences in 16S rRNA gene copy number between tissue types and animal source, except a normal distribution was specified. Likelihood ratio tests were used to determine the statistical significance of each factor (Venables & Ripley 2002). Goodness-of-fit was assessed by with a Chi-square test (Faraway 2006) and homoscedasticity was assessed by examination of residual plots. Nonparametric methods were used in univariate analyses of the metagenomic predictions because no distribution provided a reasonable fit to the data. P-values were adjusted for multiple tests using the false discovery rate (Benjamini & Hochberg 1995).

## RESULTS

### Spatial structure of the gut microbiome

We recovered 11 dominant sequence variants from field and lab-raised individuals (Fig. 2), with the remaining 749 sequence variants comprising <1% of the sequences from a given Mormon cricket. Field and laboratory-raised individuals shared 7 of the 11 sequence variants, including the most abundant *Pediococcus acidilactici* OTU that varied with mating status in a previous study (*P. acidilactici* 102222; Smith et al. 2016). The remaining five shared sequence variants were two Lactobacilliaceae (*Lactobacillus sp* and *P. acidilactici* 2), two Enterobacteriaceae (*Pantoea agglomerans* and a *Klebsiella sp*) and one Streptococcaceae (*Lactococcus garvieae*). Field-caught Mormon crickets had three taxa that were not shared with laboratory-raised individuals, while lab-raised individuals had two taxa that were not shared with field individuals (Fig. 2). Guts from two laboratory individuals were almost completely comprised of the enteric bacterium *Pantoea agglomerans* (99.3% and 80.8% of reads respectively), so we conducted our analysis with and without these individuals.

**Figure 2.**
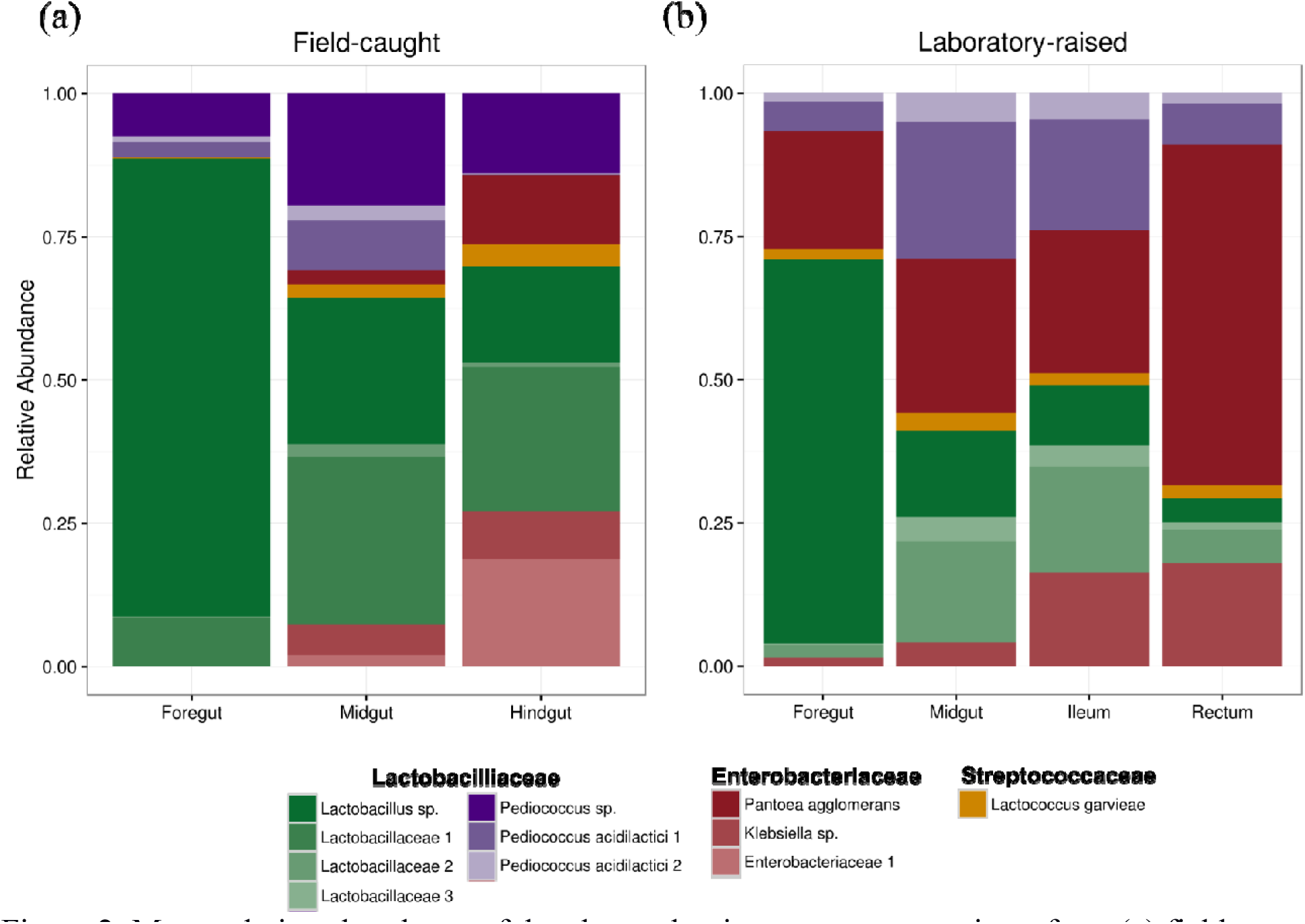
Mean relative abundance of the eleven dominant sequence variants from (a) field-caught and (b) laboratory-raised Mormon crickets from 16S rRNA Illumina sequencing.

Species richness and diversity differed among gut regions and were higher in field compared to lab-raised animals (Table 1, Fig. 3). There was no significant interaction between collection source and tissue type (Table 1), indicating that differences in alpha diversity among tissue types were shared between lab and field-caught animals. We found that the midgut was the most diverse part of the gut with two of the three measures of alpha diversity (species richness and the Chao1 diversity estimator), while the hindgut and foregut had similar levels of richness and diversity. The third metric (Shannon-Weiner) also found the foregut to be the least diverse region, but differed in that the midgut and hindgut had similar levels of species diversity (Table 1, Fig 3).

**Table 1.** Analysis of deviance comparing alpha diversity between source populations (wild or laboratory) and among tissue types (foregut, midgut, and hindgut). Values represent the F-statistic (p-value) for each term. Statistically significant terms (p<0.05) are indicated in bold. Degrees of freedom were estimated using the Kenward-Rogers approximation. In the reduced dataset, two individuals from the laboratory-reared population were removed (see Methods).

|  | Reduced Dataset |  |  | Full Dataset |  |  |
| --- | --- | --- | --- | --- | --- | --- |
|  | Species<br>Richness | Chao1 | Shannon-Weiner | Species<br>Richness | Chao1 | Shannon-Weiner |
| Source | **13.9 (0.003)** | **11.0 (0.007)** | **9.17(0.01)** | **14.0(0.002)** | **10.5 (0.006)** | **8.22 (0.013)** |
| Tissue type | **5.85 (0.010)** | **4.79 (0.02)** | **7.07 (0.005)** | **6.77 (0.004)** | **5.68 (0.008)** | **8.44 (0.001)** |
| Interaction | 0.51 (0.61) | 1.02 (0.38) | 0.98 (0.39) | 0.66 (0.53) | 1.15 (0.33) | 1.28 (0.29) |

**Figure 3.**
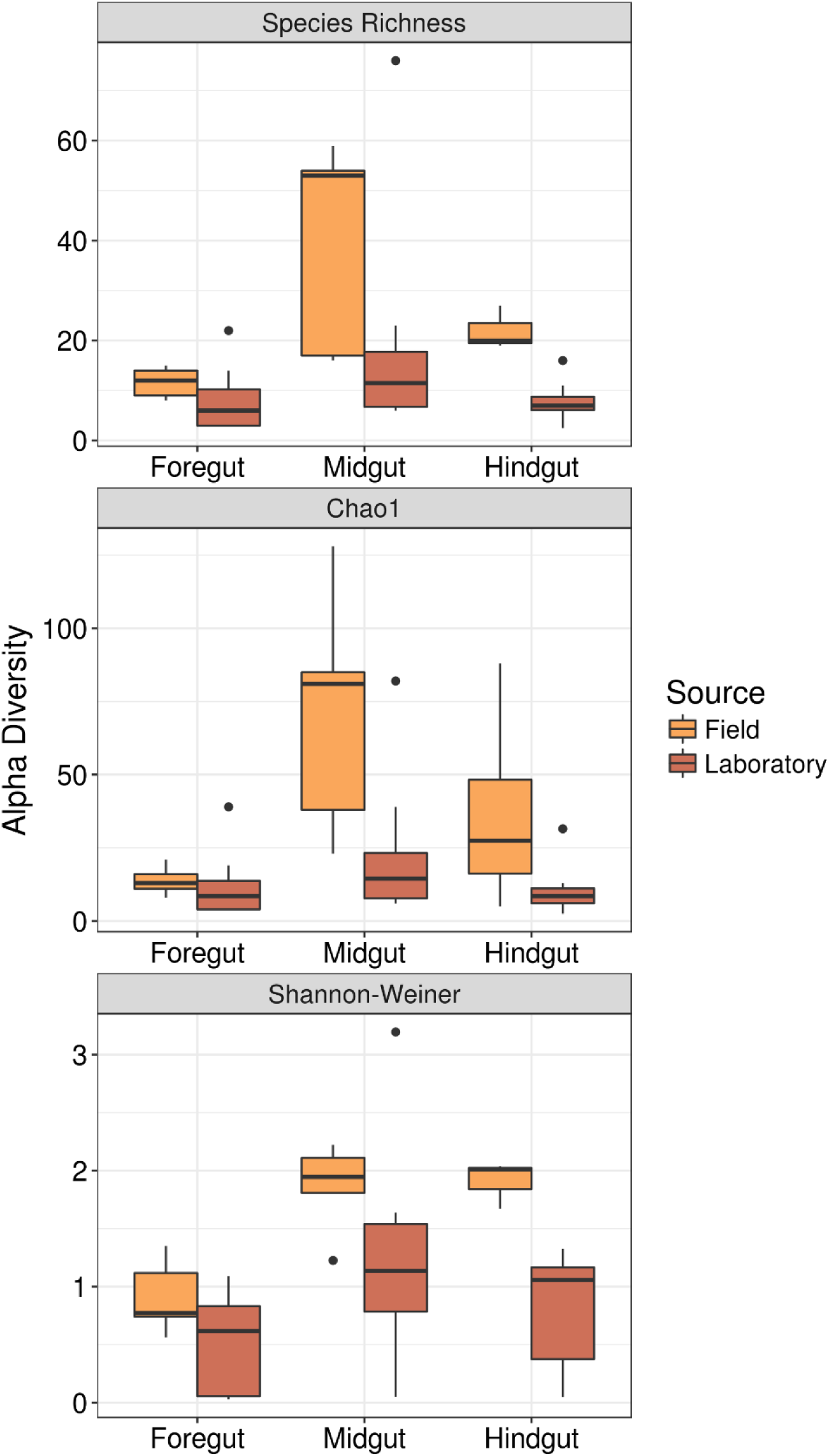
Alpha diversity in field-caught and laboratory-raised Mormon crickets.

The db-RDA analysis revealed that the structure of the gut microbiome also varied among gut regions and between field and laboratory animals (Table 2, Fig. 4a). The non-significant interaction in this analysis, however, indicates that the differences in community structure among tissue types were consistent between field and laboratory-raised individuals (Table 2). To determine which members of the gut microbiome varied among gut regions, we plotted the taxa scores from db-RDA analyses of field and laboratory Mormon crickets (Figure S1a). Three groups of bacteria appeared to separate along the gut axis: a *Lactobacillus sp.* lactic-acid bacterium associated with the foregut, *Pediococcus* lactic-acid bacteria were associated with the midgut, and *Pantoea agglomerans*, an enteric bacterium, was found in association with the hindgut. Inspection of the plots from laboratory animals, where the ileum and rectum of the hindgut were dissected separately, indicate that *P. agglomerans* is more abundant in the rectum, while the composition of the ileum, which is separated from the rectum by the colon, closely resembled that of the midgut (Figure S1b).

**Table 2.** Permutation tests of distance-based redundancy analyses of DADA2 16S rRNA sequence variants and PICRUSt metagenomic predictions. Source population (wild or laboratory) and tissue type (foregut, midgut and hindgut) were entered as factors.

|  | **Reduced Dataset** |  | **Full Dataset** |  |
| --- | --- | --- | --- | --- |
|  | *F* | *P* | *F* | *P* |
| <u>DADA2</u> |  |  |  |  |
| Source | **8.99** | **<0.001** | **7.75** | **<0.001** |
| Tissue type | **9.85** | **<0.001** | **5.49** | **<0.001** |
| Interaction | 0.26 | 0.47 | 0.52 | 0.84 |
| <u>PICRUSt</u> |  |  |  |  |
| Source | 0.23 | 0.76 | **4.31** | **0.04** |
| Tissue type | **1.61** | **0.003** | **8.09** | **0.002** |
| Interaction | 0.16 | 0.35 | 0.92 | 0.43 |

**Figure 4.**
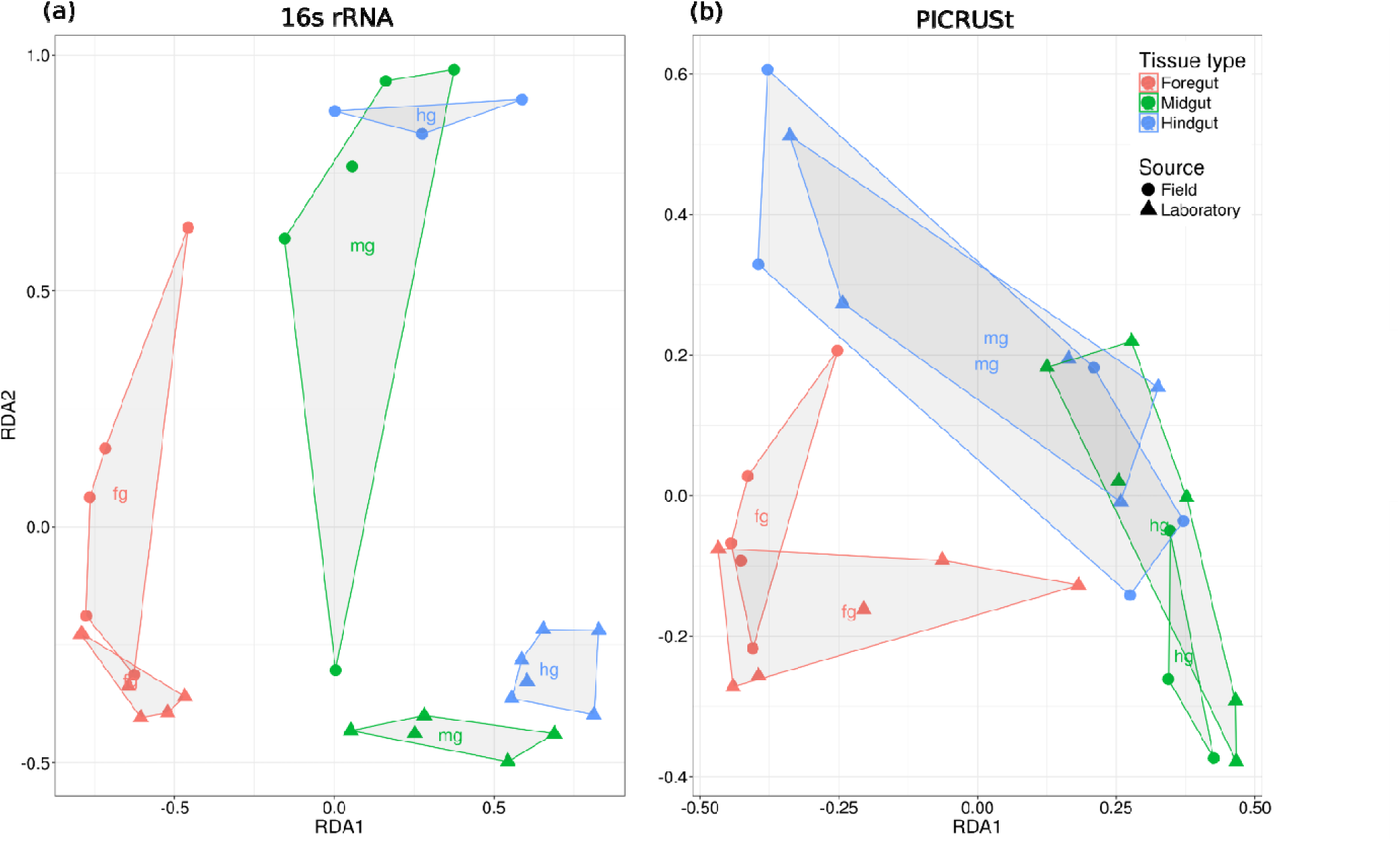
Ordination of sample scores from the db-RDA of the reduced dataset for (a) Illumina 16S sequencing and (b) PICRUSt metagenomic predictions.

Univariate analyses of these three groups largely confirmed the pattern in the ordination (Table 3, Fig. 2, Fig. S2). The interaction between tissue type and source was not significant in any of the analyses and dropped to estimate the differences in abundance between tissue types. *Lactobacillus sp*. was three times more common in the foregut than in the midgut (β=1.4 ± 0.50, p=0.02) and seven times more abundant in the foregut than in the hindgut (β=2.0 ± 0.51, p<0.001). *Pediococcus* were similar in abundance in the midgut and hindgut but 4.7 times more common in these areas than the foregut (β=1.1 ± 0.36, p=0.006). *P. agglomerans* was 209 times more abundant in the hindgut than in the foregut (β=3.8 ± 0.87, p<0.001) and twelve times more abundant in the hindgut than in the midgut (β=2.5 ± 0.82, p=0.007).

**Table 3.** Likelihood ratio tests from GLMMs fitting the abundance of sequence variants to source population (wild or laboratory) and tissue type (foregut, midgut or hindgut). Values are Chi-square (p-value). LAC1=*Lactobacillus sp.*, PED=*Pediococcus*, PAG=*Pantoea agglomerans*.

|  | Reduced Dataset |  |  | Full Dataset |  |  |
| --- | --- | --- | --- | --- | --- | --- |
|  | LAC1 | PED | PAG | LAC1 | PED | PAG |
| Source | 1.32 (0.25) | 0.80 (0.37) | 2.36 (0.12) | 0.18 (0.66) | 0.51 (0.48) | 0.01 (0.96) |
| Tissue type | **16.3 (<0.001)** | **9.25 (0.01)** | **20.7 (<0.001)** | **12.7 (0.002)** | **16.1 (<0.001)** | **41.9 (<0.001)** |
| Interaction | 0.36 (0.84) | 0.84 (0.66) | 3.86 (0.15) | 0.3 (0.86) | 0.95 (0.62) | 2.67 (0.26) |

**Table 4.** Permutation test of distance-based redundancy analysis of PICRUSt metagenomics predictions between source populations (wild or laboratory) and among tissue types (foregut, midgut and hindgut).

|  | **Reduced Dataset** |  | **Full Dataset** |  |
| --- | --- | --- | --- | --- |
|  | *F* | *P* | *F* | *P* |
| <u>Illumina 16S rRNA</u> |  |  |  |  |
| Source | **8.99** | **<0.001** | **7.75** | **<0.001** |
| Tissue type | **9.85** | **<0.001** | **5.49** | **<0.001** |
| Interaction | 0.26 | 0.47 | 0.52 | 0.84 |
| <u>KEGG Pathways</u> |  |  |  |  |
| Source | 0.23 | 0.76 | **4.31** | **0.04** |
| Tissue type | **1.61** | **0.003** | **8.09** | **0.002** |
| Interaction | 0.16 | 0.35 | 0.92 | 0.43 |

### Abundance of bacteria from 16S rRNA qPCR

The number of copies of bacterial 16S rRNA genes was significantly different among tissue types, as indicated by the significant interaction between tissue type and the source of the Mormon crickets (Analysis of deviance: Source, F_1,14_=25.9, p<0.001; tissue type, F_3,161_=7.8, p<0.001; Interaction, F_3,161_=2.8, p=0.04, Fig. 7). We decomposed the interaction to determine how the total number of 16S rRNA copies differed among tissue types within field and laboratory-raised animals. The major difference between the two sources was that in wild Mormon crickets, the midgut had the lowest abundance of all gut regions, while in laboratory-raised individuals, both the midgut and the ileum had the lowest abundance of bacterial 16S rRNA genes (Table S1, Fig. 5).

**Figure 5.**
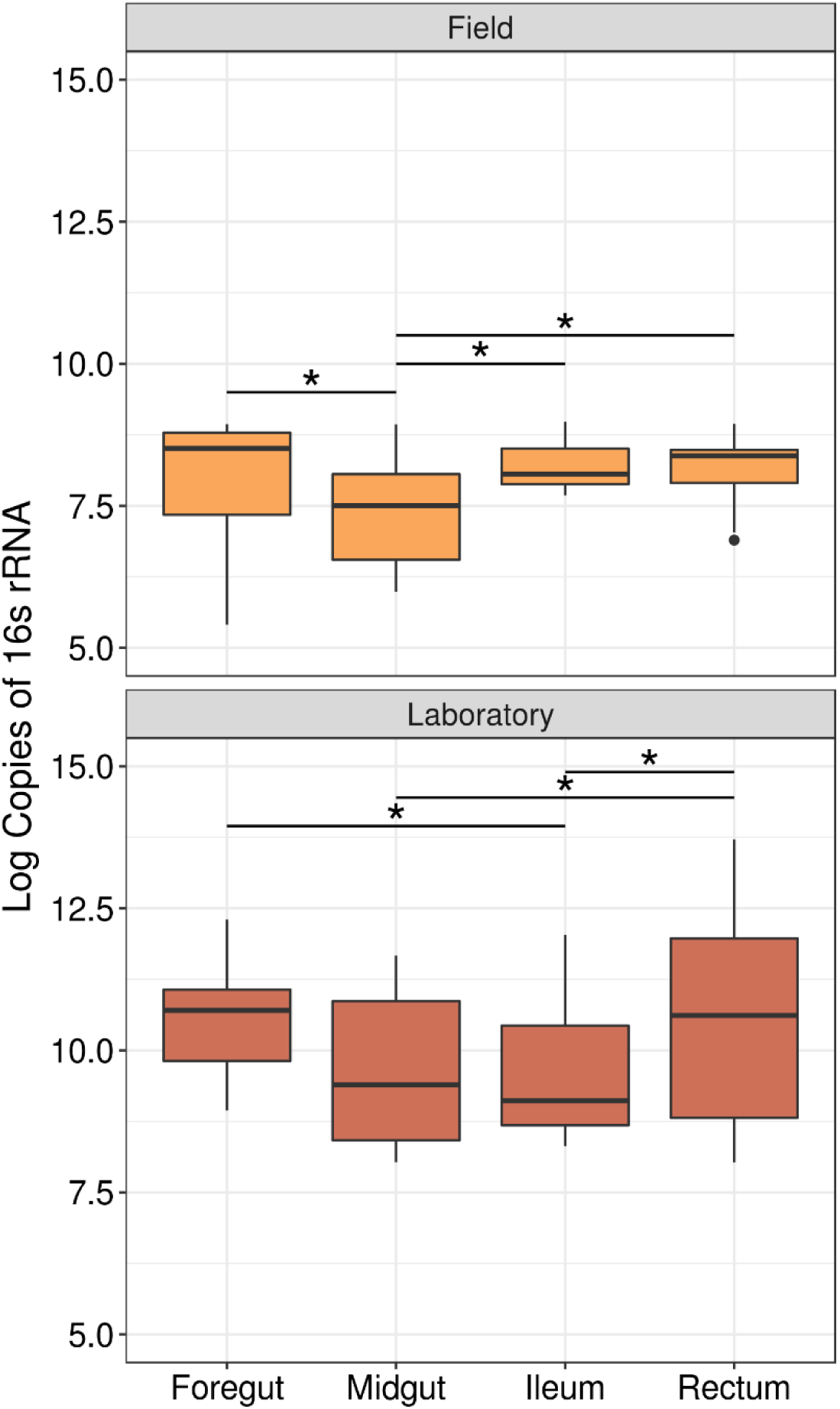
Abundance of bacterial 16S rRNA genes in the Mormon crickets gut. Bars indicate significant differences between regions (* p<0.05).

### PICRUSt metagenomic predictions

PICRUSt analysis of 16S rRNA sequence variants recovered 5,891 KEGG orthologs associated with 328 metabolic pathways. The representation of the predicted KEGG pathways differed significantly among gut regions in both the full and reduced datasets, while the source of the animals had a significant influence in the full dataset but not the reduced dataset (Table 2, Fig. 4b). Neither analysis, however, showed an interaction between tissue type and whether an animal was wild or lab-reared, indicating that metagenomic predictions differed among tissue types in similar ways (Table 2). Univariate analyses found significant differences among tissue types in most KEGG pathways (Table S2), including those that could affect host-microbe interactions via their role in nutrition, immunity, degradation of xenobiotics, and production of secondary metabolites (Fig. 6). In these functional groups, the hindgut exhibited the most abundant representation of each KEGG category, followed by the midgut and then the foregut (Fig. 6).

**Figure 6.**
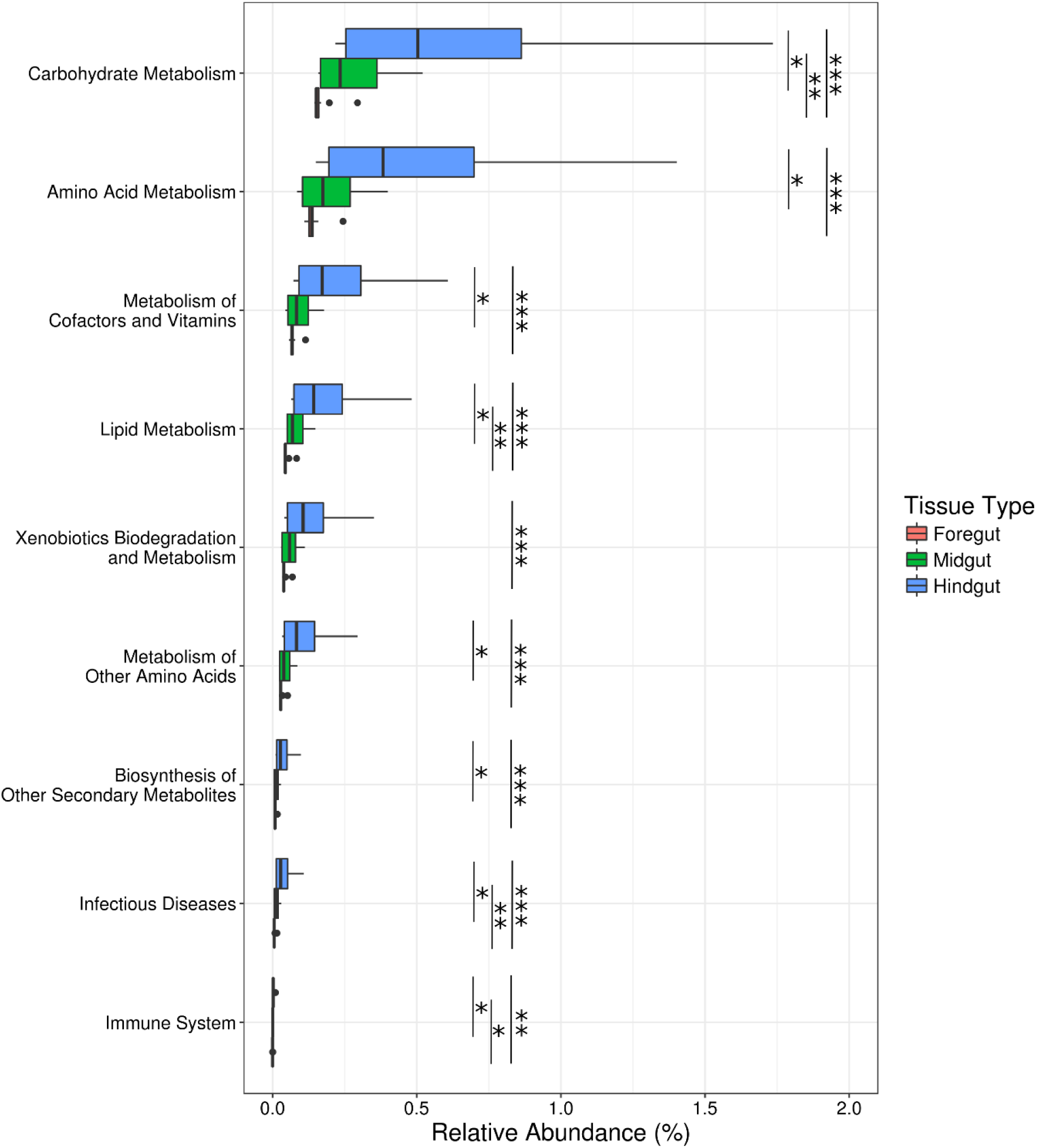
Relative abundance of KEGG pathways related to nutrition, immunity, xenobiotic degradation, and secondary metabolite production among tissue types.

### Nutrition

We searched our metagenomic predictions for specific bacterial genes known to contribute to host nutrition in orthopterans. We queried our database for enzymes capable of metabolizing the complex plant carbohydrates xylan, pectin, raffinose, and galactomannan, which are metabolized by gut bacteria in the house cricket *Achetus domesticus* (Kaufman & Klug 1991), and cellulose, an important component of the plant cell wall. We found KEGG orthologs involved in the metabolism of all these complex plant polymers, except galactomannan. The pectin metabolic pathway was also incomplete. Only pectinesterase, the first of three enzymes involved in pectin metabolism, was found among the eleven dominant taxa in Mormon crickets, although the remaining two enzymes were represented in the minority members (i.e. <1% of 16S rRNA sequences, see *Sequencing and Bionformatics*) of the gut microbiome.

The abundance of KEGG orthologs for carbohydrate metabolism in our samples were most pronounced in the hindgut (Table S3, Fig. 7a) and dominated by the enteric bacteria, particularly *Klebsiella sp*. and Enterobacteriaceae 1 (Fig 7b). Lactic-acid bacteria, however, were also represented in predictions for raffinose metabolism and enzymes capable of participating in the degradation of cellubiose to glucose via cellubiose glucohydrolase, but not in degrading cellulose to cellubiose (Fig 7b).

**Figure 7.**
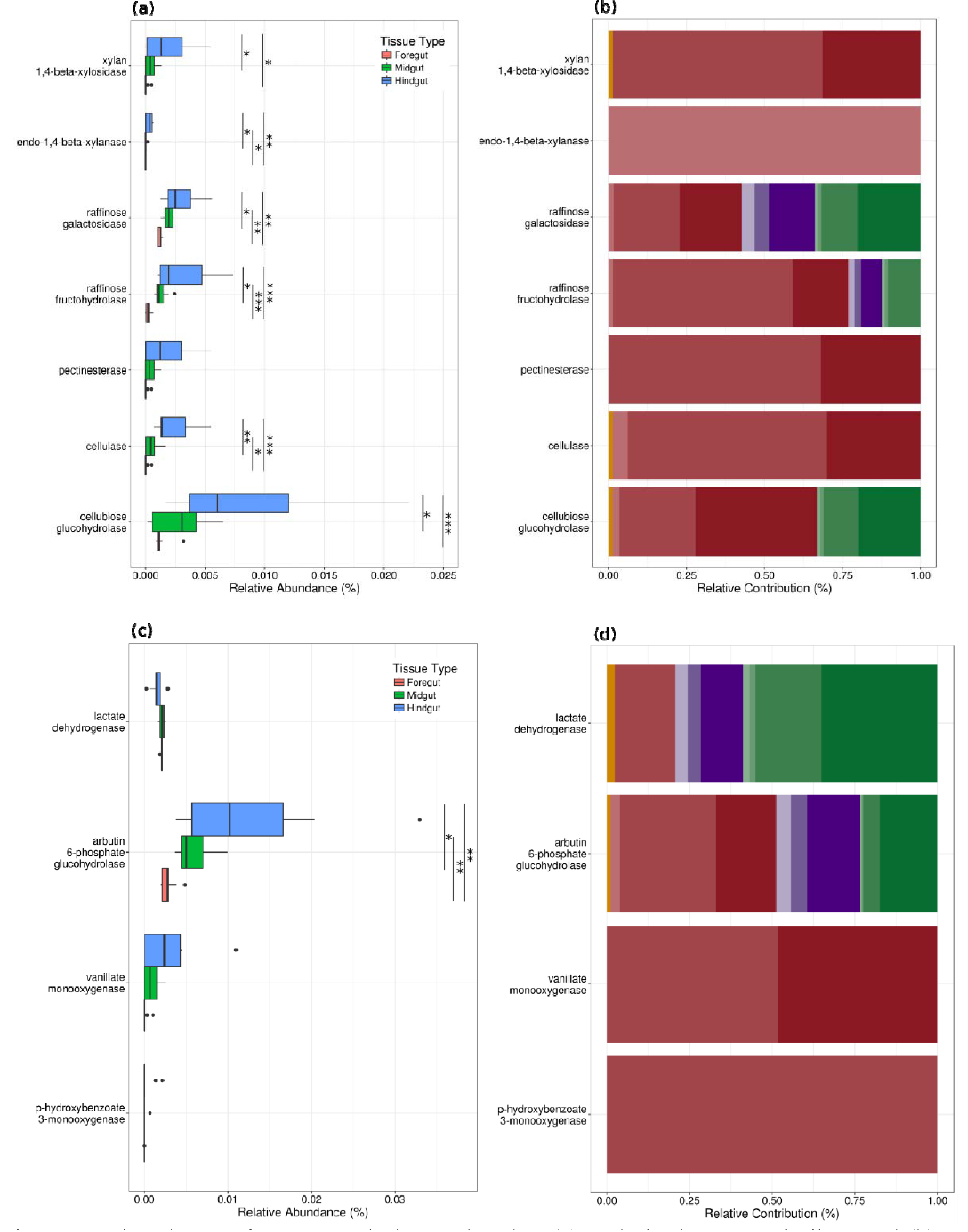
Abundance of KEGG orthologs related to (a) carbohydrate metabolism and (b) antimicrobial compound production among tissue types, and the contributions of taxonomic groups (c,d) to ortholog abundance. Key for colors representing taxonomic groups are in Figure 2.

Gut bacteria might also play a role in the production of the essential amino acid phenylalanine via the shikimate pathway, which is found in microbes and plants but not in animals (Herrmann & Weaver 1999). Phenylalanine is required for stabilization of the cuticle following molting (Bernays & Woodhead 1984) and is converted to tyrosine, the precursor of melanin, a key component of the insect immune response (González-Santoyo & Córdoba-Aguilar 2012). All enzymes in the shikimate pathway were represented in our metagenomic predictions (Fig. S3), although prephenate hydrogenase, which is required for phenylalanine synthesis, was only represented in *Lactococcus garviae. L. garviae* abundance thus might influence the availability of phenylalanine for Mormon crickets, unless they are able to acquire it in sufficient quantities directly from their diet.

### Immunity

In the locust *Schistocera gregaria* (Orthoptera), four phenols have been shown to increase resistance to microbial pathogens (Dillon & Charnley 1988, 1995): hydroquinone, 3,4-dihydroxybenzoic acid, p-hydroxybenzoic acid, and 4,5-dihydroxybenzoic acid. We found enzymes associated with the production of all these compounds except for 4,5-dihydroxybenzoic acid, which was not annotated in the KEGG database. Hydroquinone production was represented by the enzyme arbutin 6-phosphate glucohydrolase, which metabolizes arbutin, a phenolic glycoside present in leaf and fruit tissue of many plants (Xu et al. 2015).

Two enzymes were found capable of producing 3,4-dihydroxybenzoic acid. The first, vanillate monooxygenase, demethylates vanillic acid, a compound derived from lignin (Bugg et al. 2011). This is also the pathway proposed for 3,4-dihydroxybenzoic acid production in locusts based on the abundance of vanillic acid in their feces (Dillon & Charnley 1988, 1995). The second, p-hydroxybenzoate 3-monooxygenase, oxidizes p-hydroxybenzoic acid, one of the other antimicrobial phenols in locusts (Dillon & Charnley 1995). The most likely source of p-hydroxybenzoic acid in the diet of Mormon crickets is benzoic acid, which is a precursor to salicyclic acid in plants (Raskin 1992). The enzyme responsible for catalyzing the conversion of benzoic acid to p-hydroxybenzoic acid (benzoate 4-monooxygenase), however, was not found among the 11 dominant taxa in our samples, although it was present in the minority members of the Mormon cricket gut microbiome. Production of p-hydroxybenzoic acid in appreciable concentrations is thus less likely than for hydroquinone or 3,4-dihydroxybenzoic acid.

Like carbohydrate metabolism, the hindgut (Fig. 7c) and enteric bacteria (Fig. 7d) dominated the abundance of KEGG orthologs implicated in the production of antimicrobial phenols in our samples, with the exception of hydroquinone, which was represented to varying degrees among the lactic-acid bacteria. Notably, *P. agglomerans,* which has been reported to participate in the production of 3,4-dihydroxybenzoic acid in locusts (Dillon & Charnley 1995), was not among taxa responsible for the occurrence of vanillate monooxygenase in our samples (Fig 7d).

Finally, we searched for three other known contributors to pathogen defense: bacterocins, antibiotics, and lactate dehydrogenase, which provides protection from pathogens in the gut by reducing pH (Servin 2004). We found lactate dehydrogenase to be equally represented among gut regions (Fig 7c), and lactic-acid bacteria were the main contributors to our samples (Fig. 7d). We found three bacteriocins in the KEGG database: nisin, mutacin, and *blp*-derived bacterocins. None of these were found in our metagenomics predictions, perhaps not surprising considering their association with *Streptococcus,* which was not among the top 11 taxa in our samples (Fig. 2). The bacteriocins we would expect to find based on taxonomy (e.g. pediocin for *Pediococcus*) were not annotated in the KEGG database.

Turning to the antibiotics, we found enzymes involved in the production of streptomycin, penicillin, and novobiocin, but not all enzymes required for their synthesis were present (data not shown). We did find β-lactamase, which confers resistance to β-lactam antibiotics (e.g. penicillins, cephalosporins, monobactams, and carbapenems; Drawz & Bonomo 2010), represented among all the Enterobacteriaceae and *Pediococcus* taxa in our samples, but not among *Lactobacillis sp.* other Lactobacillaceae (data not shown). This suggests that lactate production and antibiotic resistance could play a role in microbe-microbe interactions in the Mormon cricket gut microbiome.

### Phylogenetic analysis of cultured isolates

Thirteen strains were cultured from the Mormon cricket gut based on 99% sequence similarity of their near full-length 16S rRNA genes (mean ± sd: 1406 ± 30bp). Six were lactic-acid bacteria (Lactobacillaceae) and seven were enteric bacteria (Enterobacteriaceae).

The lactic-acid bacteria fell into two clades in our phylogenetic analysis (Fig. 8). The first clade was comprised of *Pediococcus acidilactici* isolates derived from environmental sources, such as plants and various human foodstuffs, as well as strains from the human gut. Similarity to sequences from the BLAST search was high (>99.5%) and branch lengths were short, indicating that *Pediococcus* from the Mormon cricket gut are not highly derived from their relatives, as has been found for *Lactobacillus* species isolated from bees (Fig. 8; McFrederick et al. 2013).

**Fig 8.**
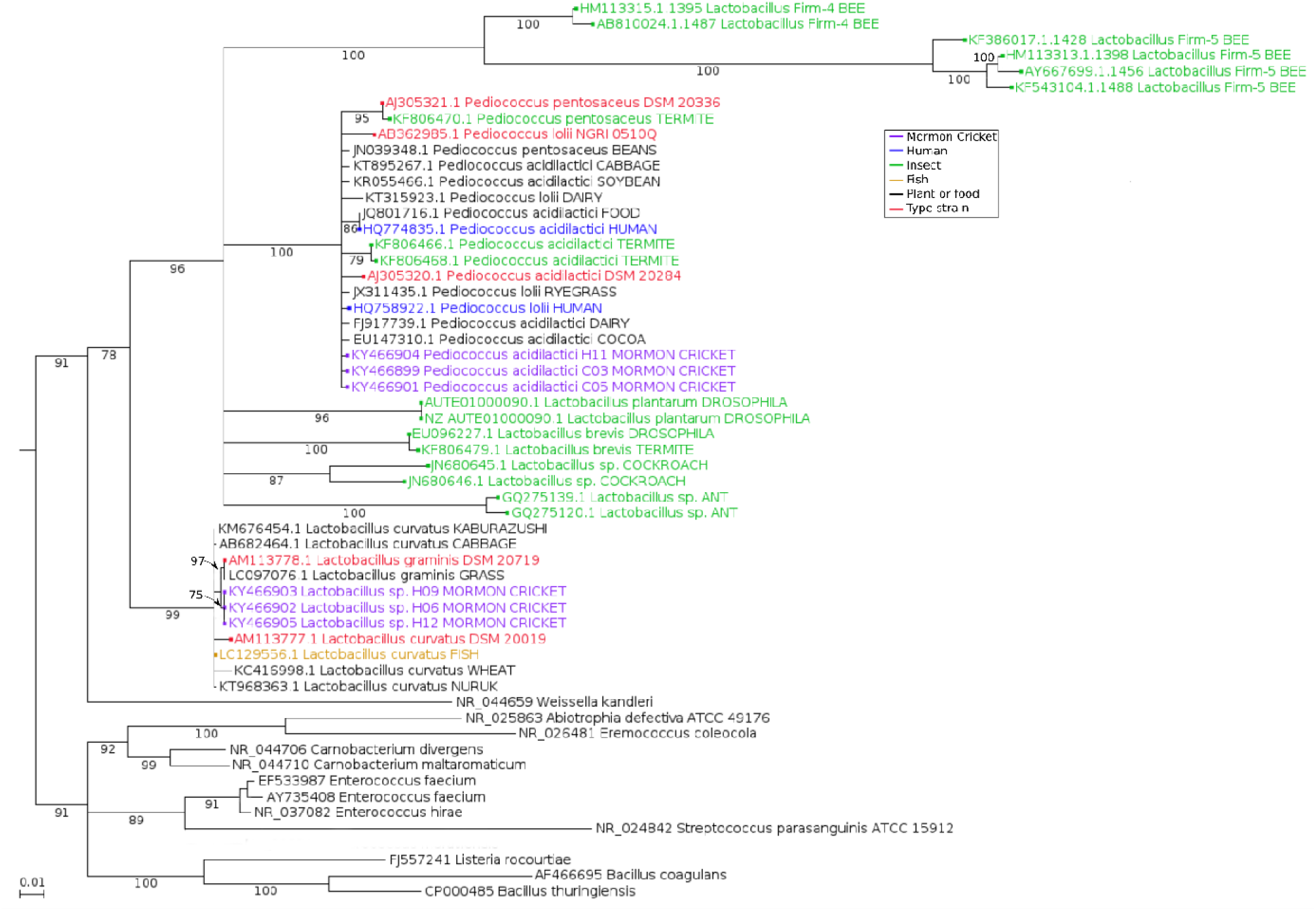
Maximum likelihood estimation of phylogenetic relationships among lactic-acid bacteria 16S rRNA sequences from Mormon cricket gut isolates and their relatives. Branches with bootstrap support <75% are collapsed.

Our search for *Pediococcus* sequences from insect guts in Genbank recovered sequences from the termites *Macrotermes bellicosus* and *M. subhyalinus*, which formed their own well-supported clade (Fig 8). Cultured *Pediococcus acidilactici* shared 100% sequence identity in the V4 region with the *P. acidilactici 1* phylotype sequenced using the Illumina platform in this study and with the *P. acidilactici* (102222) phylotype associated with variation in mating status in Mormon crickets (Smith et al. 2016). Morphologically, *Pediococcus acidilactici* were nonmotile and spherical (0.8 – 1.0 μm), often dividing to form pairs as described for other *Pediococcus*. As other members of the genus, the *P. acidilactici* were gram-positive, non-motile, facultatively anaerobic, grow at low pH, and produce lactate from lactose (Table S2).

The second clade of lactic-acid bacteria was comprised primarily of plant-associated *Lactobacillus.* Unlike *P. acidilactici*, these *Lactobacillus* formed a distinct clade with good branch support (Fig 8), indicating it is genetically distinct enough at the 16S rRNA locus to distinguish itself from other clades in the phylogeny. Similar to *P. acidilacitici*, these *Lactobacillus* had high sequence similarity (>99.5%) to other members of the clade and a short branch length, indicating that while it is distinct enough to form its own clade, it is not highly derived from its relatives at the 16S rRNA locus.

Our Genbank search for *Lactobacillus* isolated from insect guts found sequences from ants, bees, and termites, and fruit flies, all of which fell into a different clade than *Lactobacillus* isolated from Mormon crickets. *Lactobacillus* from these taxa thus appear to have a different evolutionary history. *Lactobacillus* isolates shared 100% sequence identity in the V4 region with the Lactobacillaceae 2 phylotype sequenced using the Illumina platform in this study. Morphologically, these *Lactobacillus* appear as non-motile straight rods, approximately 1.3-2 μm in length and 0.8-1.0 μm wide and are gram-positive, non-motile, facultatively anaerobic, grow at low pH, and produce lactate from lactose (Table S2).

The seven Enterobacteriaceae strains were most similar to *Enterobacter* strains in our BLAST search, which recovered sequences from a variety of plant and animal sources (sequence similarity=98.7-99.8%). Our survey of Genbank found *Enterobacter* from alimentary tracts of a diverse group of insects, including termites, cockroaches, flies, beetles, stink bugs, bees, ants, and moths. Like other studies (Brenner et al. 2005), however, the 16S rRNA gene did not have enough signal to resolve relationships among *Enterobacter* and its relatives (data not shown) so we present a simpler phylogeny with the Mormon cricket isolates and type strains from the family (Fig. 9).

**Figure 9.**
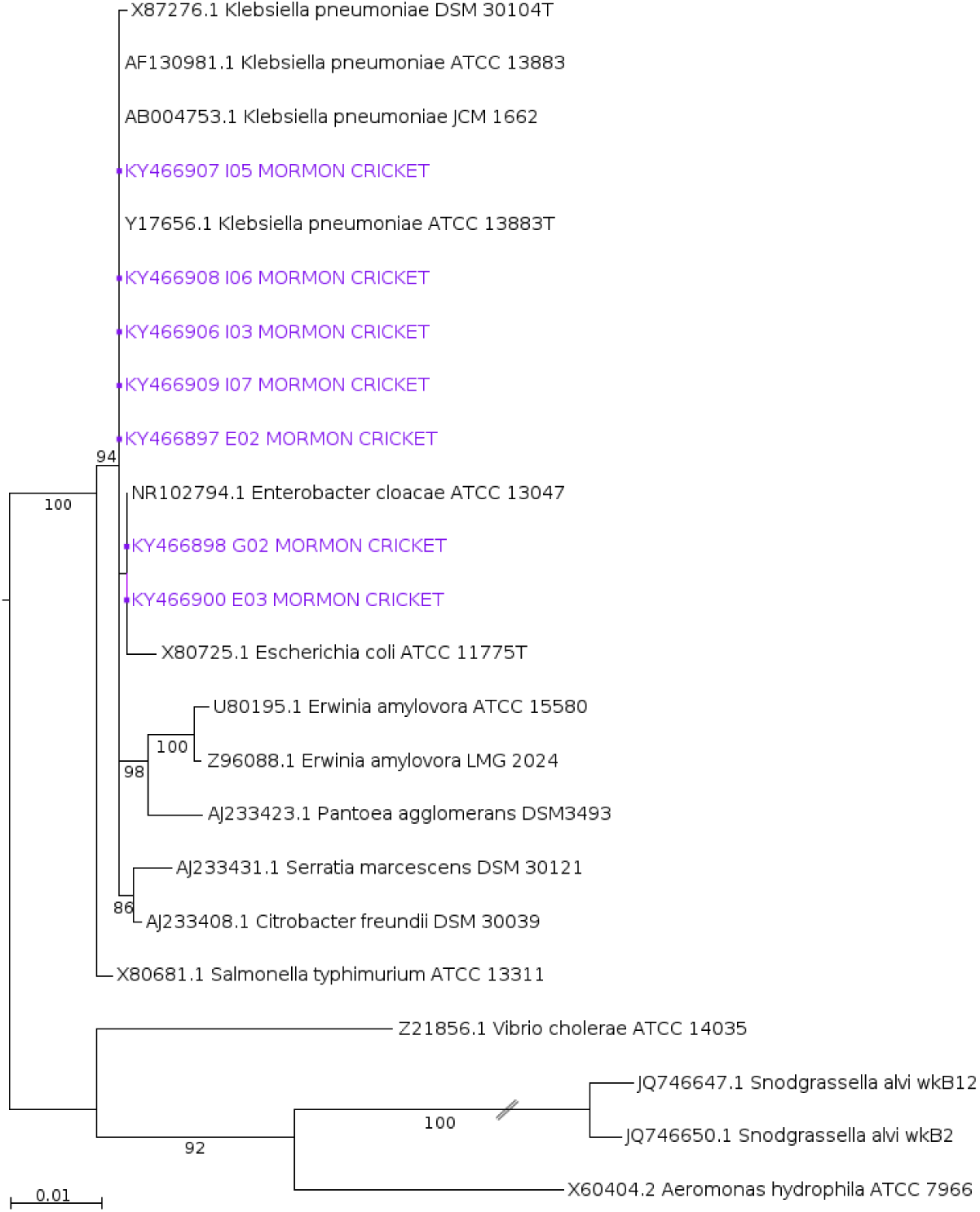
Maximum likelihood estimation of phylogenetic relationships among enteric bacteria 16S rRNA sequences from Mormon cricket gut isolates and type strains. Branches with bootstrap support <50% are collapsed.

We found that our Mormon cricket isolates were interspersed with *Enterobacter*, *Klebsiella*, and *Escherichia* type strains. A multilocus sequencing approach is thus needed to improve the inference (Brenner et al. 2005). All seven strains isolated from Mormon crickets had 100% identity at the V4 region with the *Klebsiella* phylotype sequenced on the Illumina platform, however the phylogenetic (Fig. 9) and phenotypic data (Table S2) suggest that *Klebsiella* is unlikely to be a correct taxonomic assignment. Unlike most *Klebsiella*, cultured strains were motile, which is more typical of *Enterobacter* and other Enterobacteriaceae (Brenner et al. 2005). Morphologically, all isolates were straight rods, approximately 0.8-1.0 μm in length and 0.6-0.8 μm wide. Strains were gram-negative and facultatively anaerobic (Table S2).

## DISCUSSION

We found striking differences in the diversity and structure of the gut microbiome in the Mormon cricket *Anabrus simplex*. While most taxa were represented in the foregut, midgut and hindgut, there were dramatic differences in relative abundance within the Lactobacillaceae and between the Lactobacillaceae and Enterobacteriaceae, the main families recovered in our culture and culture-independent studies. Predictions of their metabolic capabilities using PICRUSt suggest the potential for these gut bacteria to participate in the metabolism of complex carbohydrates and defense against microbial pathogens, particularly among the enteric bacteria in the midgut and hindgut, and to a lesser extent, the lactic-acid bacteria. Finally, our phylogenetic analysis of cultured isolates found that Mormon cricket gut bacteria are not highly derived from related bacteria associated with plants or the guts of other animals, suggesting that gut bacteria are either acquired from the environment in each generation or have not been restricted to Mormon crickets over appreciable periods of evolutionary time. Our findings have important implications for our understanding of the ecological and evolutionary processes that influence the assembly and function of gut microbial communities, as it suggests that host-microbe and microbe-microbe interactions shape the abundance and distribution of the gut microbiome.

Our finding that bacterial abundance is lower in the midgut is in agreement with reports from other orthopterans (Ulrich, Buthala & Klug 1981; Hunt & Charnley 1981) and insects (Köhler et al. 2012), and has been attributed to characteristics that make the midgut less hospitable to bacteria than other regions of the alimentary tract (Douglas 2015). The midgut in insects secretes a host of digestive enzymes, is immunologically active, and lined by the peritrophic membrane, which acts as a protective barrier that restricts microbes to the lumen and protects the epithelium (Douglas 2015). In the two orthopterans that have been studied in detail, bacteria are found in the midgut lumen but not in association with the epithelium (Hunt & Charnley 1981; Mead, Khachatourians & Jones 1988). As a consequence, midgut bacteria might need to be continually replenished from ingested food (Blum et al. 2013) because the peritrophic membrane is continually shed into the hindgut. In some insects, specialized midgut crypts provide niches that microbes colonize (Kikuchi et al. 2005; Bistolas et al. 2014), however we did not observe analogous structures in Mormon crickets (Fig. 1).

The midgut is particularly vulnerable to pathogens because the lack of an endocuticle leaves the epithelium exposed once the peritrophic membrane is penetrated (Lehane & Billingsley 1996). The Mormon cricket midgut was populated by lactic-acid bacteria, with *Pediococcus* specifically exhibiting greater abundance in the midgut (and hindgut) than in the foregut. Lactic-acid bacteria are known for their beneficial effects in insects, increasing resistance to parasites in bees (Forsgren et al. 2010) and promoting development in fruit flies by enhancing proteolytic activity (Erkosar et al. 2015) and upregulating host ecdysone and insulin-like peptides (Storelli et al. 2011). Lactic-acid bacteria are also known to suppress pathogenic bacteria by reducing pH through the production of lactate and by producing a number of antimicrobial compounds, such as hydrogen peroxide and bacteriocins (Cintas et al. 2001).

A previous study found that sexual interactions in Mormon crickets influences the abundance of three *Pediococcus* phylotypes (Smith et al. 2016), however spatial information on where in the gut *Pediococcus* is located has been unavailable until now. *Pediococcus* in the midgut could provide immunological or nutritional benefits to Mormon crickets, as has been shown for *P. acidilactici* in other animals (Castex et al. 2008, 2009). We found that the capacity for lactate production in our samples was dominated by *Pediococcus* and other lactic-acid bacteria, although the abundance of the enzyme mediating lactate production was not higher in the midgut relative to other regions based on our metagenomics predictions. The cultured isolates of *P. acidilactici* obtained from Mormon crickets in this study will enable future experimental and comparative genomic approaches to evaluate these hypotheses.

Lactic-acid bacteria were also common in the foregut, which was dominated by a *Lactobacillus* that averaged 73.9% of the sequences recovered from this region. Bignell (1984) noted that the foregut of insects tends to be the most acidic compartment, however studies that measure the physiochemical environment and characterize microbiome composition of the foregut are rare (but see Köhler et al. 2012). This is because the endocuticle, lack of differentiated cells for absorption of nutrients, and frequent purging of consumed material into the midgut provides little opportunity for foregut microbes to contribute to host nutrition. The large differences in community structure between the foregut and the rest of the alimentary tract in our study does illustrate the dramatic transition in microbial communities between what is ingested and what can colonize the more distal regions of the gut. Our metagenomics predictions also suggest that the foregut is not the site of extensive carbohydrate metabolism or pathogen defense for most of the pathways we examined.

In contrast to the foregut and midgut, the hindgut was characterized by a dramatic increase in enteric bacteria (Enterobacteriaceae). Ordination of the laboratory Mormon cricket samples indicated that the rectum, not the ileum, was primarily responsible for the difference in community structure in the hindgut. Enterobacteriaceae comprised 83.5% of the sequences from the rectum compared to 57.5% from the ileum in laboratory-raised animals, which was more similar to the midgut in community structure (Fig. 4a). This distinction is potentially important because higher digestive efficiency in conventional compared to germ-free crickets has been attributed to microbial colonization of the ileum in the orthopteran *A domesticus* (Kaufman & Klug 1991).

Metabolism of the specific complex carbohydrates attributed to bacteria in this study were also identified in our metagenomic predictions and localized to the hindgut, as well as enzymes involved in the production of the essential amino acid phenylalanine via the shikimate pathway. Phenylalanine is a precursor for tyrosine, which is required to stabilize the cuticle during molting (Bernays & Woodhead 1984) and in phenoloxidase synthesis, an important component of the insect immune system (González-Santoyo & Córdoba-Aguilar 2012). Tradeoffs between allocation of tyrosine to immune function and cuticle formation during development in Mormon crickets (Srygley 2012) might be impacted by microbial contributions to amino acid production if sufficient quantities of these amino acids are not obtained directly from their diet.

Of the three enteric bacteria represented in this study, *P agglomerans* was common to both field and lab individuals and increased in abundance in the hindgut. *Pantoea* are known plant pathogens and have been associated with a variety of medical conditions in humans (Walterson & Stavrinides 2015). In insects, however, *Pantoea* have been shown to have mutualistic associations with their host. They are required for the completion of development in stinkbugs (Hosokawa et al. 2016; but see Dillon & Charnley 2002), produce compounds that attract insects to their host plants in flies (Robacker, Lauzon & He 2004; Maccollom et al. 2009), and in the orthopteran *Schistocerca gregaria*, produce a key component of the locust aggregation pheromone (Dillon, Vennard & Charnley 2000, 2002) and reduce susceptibility to microbial pathogens through the production of phenols (Dillon & Charnley 1986, 1995).

Our metagenomics predictions suggest that enteric bacteria in Mormon crickets might be capable of producing at least two of the antimicrobial phenols identified in *S. gregaria*, although *P. agglomerans* was not identified as an important contributor in our study. This illustrates a limitation of PICRUSt, as genomes in the IMG database used to make inferences about gene content may miss important among-strain variation in metabolic capabilities. *P. agglomerans* derived from *S. gregaria* are likely to have acquired this capability independently, unless the metabolic pathway is different from the one analyzed here or the taxonomic designation reported by Dillon and Charnley (1986, 1995) is incorrect.

Metagenomic analyses are also dependent upon annotation of the relevant pathways in the KEGG database. We were unable, for example, to assess the potential for the Mormon crickets microbiome to produce bacteriocins or the aggregation pheromone guaiacol, a bacterial metabolite produced by *P. agglomerans* in *S. gregaria* (Dillon et al. 2000, 2002), because the predicted pathways are not annotated in the KEGG database. The role of the gut microbiome in protecting Mormon crickets from their own pathogens (MacVean & Capinera 1991) and its influence on aggregation behavior (MacVean 1987; Simpson et al. 2006) is thus an important direction for future research.

## Conclusion

Variation in morphology and physiology is thought to differentiate niches within the gut that influence the organization of the microbiome. Our study describes at high resolution how bacterial communities vary among gut regions, and suggests that host-microbe and/or microbe-microbe interactions have a role in how microbial communities are assembled and maintained. While the metagenomics predictions gleaned from our study suggests that some of these bacteria might benefit Mormon cricket nutrition, immunity, and perhaps even modulate social behavior, experiments are needed to evaluate this possibility. Our establishment of methods for culturing Mormon cricket gut bacteria will enable experimental and comparative genomic approaches in the future to infer the ecological and evolutionary consequences of host-microbe symbiosis.

## Data Accessibility

Sequences are archived under NCBI BioProject PRJNA362233.

## Acknowledgements

We thank Laura Senior, USDA-ARS, for help with field collections and rearing laboratory insects. We also thank Bruce Shambaugh and the Cheyenne office of USDA-APHIS, PPQ for assisting with field collections, and Nancy Moran, UT-Austin, for providing the standard for the qPCR experiment. Work was funded by National Science Foundation award DEB-1354666 and the W.M. Wheeler Lost Pines Endowment from the University of Texas at Austin to UGM.

